# Identification of new therapeutic targets in *CRLF2-*overexpressing B-ALL through discovery of TF-gene regulatory interactions

**DOI:** 10.1101/654418

**Authors:** Sana Badri, Beth Carella, Priscillia Lhoumaud, Dayanne M. Castro, Claudia Skok Gibbs, Ramya Raviram, Sonali Narang, Nikki Evensen, Aaron Watters, William Carroll, Richard Bonneau, Jane A. Skok

## Abstract

Although genetic alterations are initial drivers of disease, aberrantly activated transcriptional regulatory programs are often responsible for the maintenance and progression of cancer. *CRLF2*-overexpression in B-ALL patients leads to activation of JAK-STAT, PI3K and ERK/MAPK signaling pathways and is associated with poor outcome. Although inhibitors of these pathways are available, there remains the issue of treatment-associated toxicities, thus it is important to identify new therapeutic targets. Using a network inference approach, we reconstructed a B-ALL specific transcriptional regulatory network to evaluate the impact of *CRLF2*-overexpression on downstream regulatory interactions.

Comparing RNA-seq from *CRLF2*-High and other B-ALL patients (*CRLF2*-Low), we defined a *CRLF2*-High gene signature. Patient-specific chromatin accessibility was interrogated to identify altered putative regulatory elements that could be linked to transcriptional changes. To delineate these regulatory interactions, a B-ALL cancer-specific regulatory network was inferred using 868 B-ALL patient samples from the NCI TARGET database coupled with priors generated from ATAC-seq peak TF-motif analysis. CRISPRi, siRNA knockdown and ChIP-seq of nine TFs involved in the inferred network were analyzed to validate predicted TF-gene regulatory interactions.

In this study, a B-ALL specific regulatory network was constructed using ATAC-seq derived priors. Inferred interactions were used to identify differential patient-specific transcription factor activities predicted to control *CRLF2*-High deregulated genes, thereby enabling identification of new potential therapeutic targets.

## INTRODUCTION

Acute lymphoblastic leukemia (ALL) is the most common cancer in children. Historically, clinical criteria have driven risk stratification for these patients, however over time many genetic alterations have been identified as prognostic predictors. Several groups have described varied genetic signatures associated with pediatric ALL with the aim of elucidating their contributions to leukemogenesis^1, 2^. Improved risk stratification based on genetic signature has altered treatment and led to significant improvements in overall survival^3^. However, about 20% of patients fail current treatment strategies or die following relapse. Moreover, adults with ALL have a worse prognosis and an average overall survival of 35-50%^3^. Therefore, it is important to gain a better understanding of the mechanisms by which genetic alterations drive leukemogenesis to refine therapies that target the disease-essential pathways involved^4^.

Genetic alterations that lead to overexpression of the cytokine receptor-like factor 2 (*CRLF2*) gene have been associated with a high-risk subset of pediatric patients with B-cell acute lymphoblastic leukemia (B-ALL). The *CRLF2* gene encodes the thymic stromal lymphopoietin receptor (TSLPR) which forms a heterodimer with IL-7 receptor alpha (IL7RA) to bind TSLP^5^. Binding of TSLP to the IL7RA-TSLPR complex signals the phosphorylation of Janus kinase 1 (JAK1) and Janus kinase 2 (JAK2), leading to the activation of the JAK-STAT signaling pathway^6^. Studies have shown that stimulation of TSLP in B-ALL not only induces activation of the JAK-STAT pathway, but also activates the PI3K/mTOR^7^ and ERK/MAPK signaling pathways^3^.

*CRLF2* overexpression occurs in 5-15% of patients with B-ALL and in 50-60% of pediatric B-ALL patients with Down syndrome (DS)^6, 8^. Overexpression can occur either from a chromosomal translocation between *CRLF2* and the immunoglobulin heavy chain locus (*IGH*) on chromosome 14 (*IGH-CRLF2*) or from an interstitial deletion of the pseudoautosomal region of the X/Y chromosomes resulting in the fusion of *CRLF2* to the *P2RY8* gene (*P2RY8-CRLF2*)^9^ as shown in **Figure 1a**. *CRLF2* deregulation very rarely occurs via activating mutations of *CRLF2*. *IGH-CRLF2* translocation occurs in precursor cells and is thought to result from aberrant rejoining during V(D)J-recombination^10, 11^. This translocation is most commonly found in adolescents and adult patients and is typically associated with a poor prognoses, while the *P2RY8-CRLF2* fusion is found in younger patients^3^.

**Figure 1:**
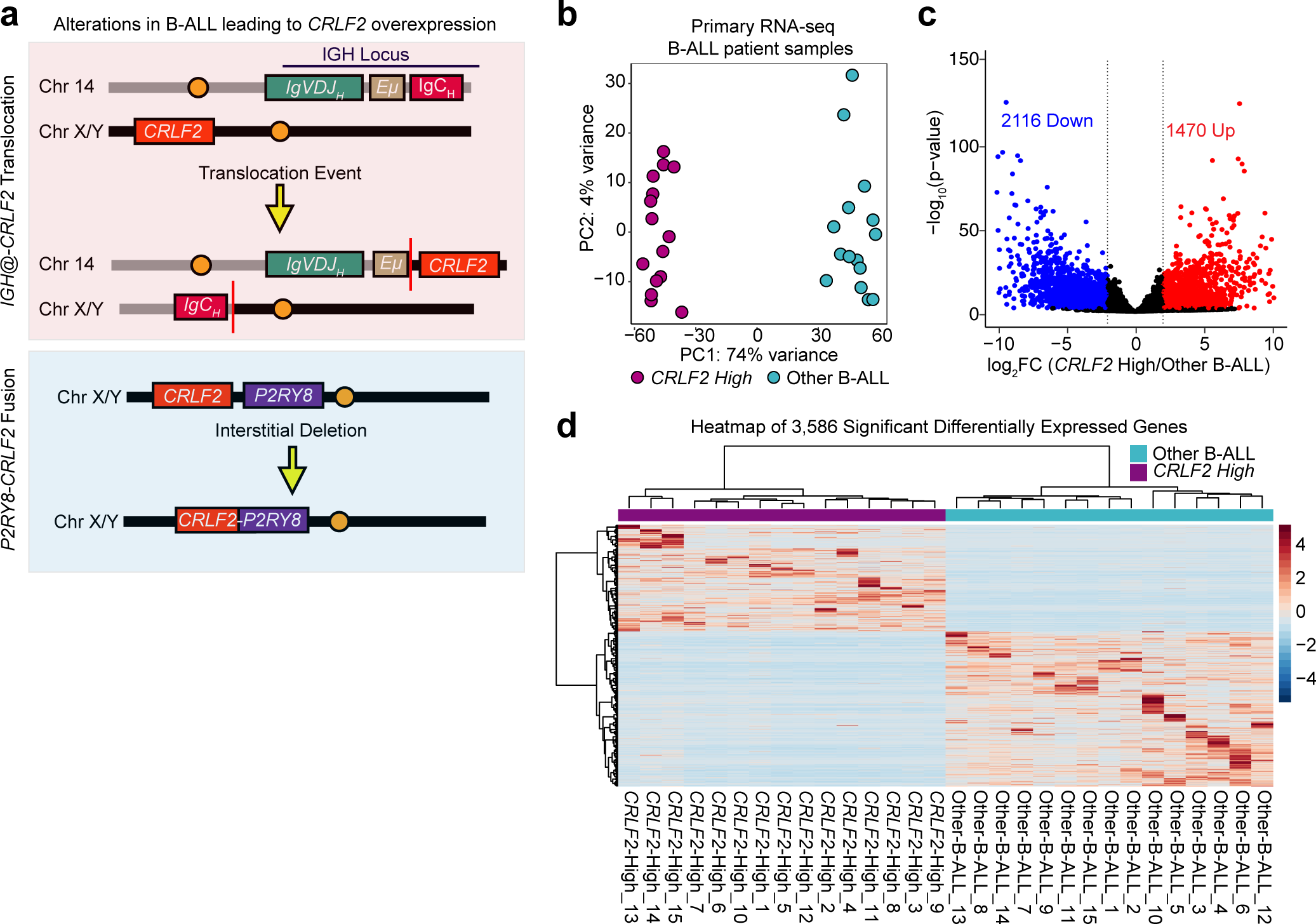
Genome-wide transcriptional changes in CRLF2-overexpressing primary patient samples. (a) Schematic of chromosomal alterations, IGH-CRLF2 (top) and P2RY2-CRLF2 (bottom) leading to CRLF2 overexpression. (b) PCA analysis of 15 CRLF2-High (purple, n=15) and 15 Other B-ALL (teal, n=15) patient RNA-seq samples. (c) Volcano plot of the differentially expressed genes (DESeq2: FDR=1%, |log2 fold change| >2) between CRLF2-High and other B-ALL patients. Red and blue points correspond to up and down-regulated genes (n=1470 and 2116, respectively) in CRLF2- High samples. (d) Expression heatmap of 3,586 differentially expressed genes across patient samples.

*CRLF2* chromosomal alterations are often accompanied by mutations in *JAK1*, *JAK2* and *Ikaros* (*IKZF1*) genes. Several groups have suggested that aberrant *CRLF2* signaling cooperates with mutant JAK and IKZF1 activity to promote the development of leukemia^9, 12, 13^.

As a result, the focus has shifted towards the use of signal transduction inhibitors (STIs) to target JAK-STAT, PI3K and MAPK signaling pathways^3^. Although, STIs have shown promise in early clinical trials^14–16^, broad application of signal transduction inhibitors is challenging due to the interconnected roles of their targets in biological processes (i.e. JAK kinases) including immunity and hematopoiesis^17^. Moreover, it has been shown that mutated *JAK2* is required for the initiation of leukemia, but it is not necessary for its maintenance ^18^.

More recently, studies have constructed short and long chimeric antigen receptor (CAR)- expressing T-cells to target *CRLF2* receptor and have demonstrated only the short CARs targeting *CRLF2* can eliminate the leukemia in cell lines and xenograft models of *CRLF2*- overexpressing ALL^19^. *CRLF2*-targeted CAR T cells represent a powerful immunotherapeutic strategy for this cohort of B-ALL patients, nonetheless there are severe toxicity issues associated with this type of therapy, related to damage of normal tissues that express this antigen^20^.

Exome and targeted sequencing of DS-ALL with *CRLF2* rearrangements has uncovered driver mutations in *KRAS* and *NRAS* that are mutually exclusive to *JAK* mutations. *RAS* and *JAK* mutations appear to occur at a similar frequency. Analysis of clonal architecture demonstrates that the majority of clones initiated by *CRLF2* rearrangements are expanded into distinct sub-clones by both *RAS* and *JAK2* mutations. Interestingly, the majority of relapsed cases switch from a *JAK2*-mutated sub-clone to a *RAS*-mutated sub-clone, indicating there are alternate drivers in relapsed *CRLF2*-overexpressing DS-ALL^21^. Although, this particular study focused on DS-ALL, there is evidence that alternate genes can be responsible for the maintenance of *CRLF2*-overexpressing cells^18^, therefore, there is a need to more closely investigate the impact of the altered *CRLF2* in B-ALL downstream of oncogenic signaling, to identify alternate targets that can be evaluated for therapy.

To address the issue of alternate targets, we first sought to identify transcriptional regulators that control differentially expressed genes associated with *CRLF2*-overexpression through a comparative analysis of *CRLF2*-High cells versus other B-ALL samples that express *CRLF2* at low levels in primary patient samples. Differential analysis of RNA-seq samples resulted in robust genome-wide gene expression changes, with an expected significant enrichment in cytokine-mediated signaling pathways. Using ATAC-seq data, patient-specific chromatin accessibility landscapes were defined and interrogated to identify altered regions that could be linked to transcriptional changes. While strong changes in accessibility were detected in patients, only a proportion of these were linked to transcriptional changes. As a result, we hypothesized that differential binding of TFs at ATAC-seq peaks are influencing gene expression changes.

Using 868 B-ALL patient samples from the NCI TARGET database, we constructed A B- ALL transcriptional network to define the interactions between transcription factors (TFs) and the genes they regulate^22–24^. With RNA-seq, ATAC-seq, and TF-motif analysis, we inferred the targets of TFs linked to the gene set associated with the *CRLF2*-High cohort using the Inferelator algorithm^25, 26^. The network was then used to predict differential transcription factor activity (TFA) in *CRLF2*-High versus other B-ALL (*CRLF2*-Low) patient samples. This approach enabled the identification of a *CRLF2*-High associated sub-network consisting of thirty-four potential regulators of differentially expressed genes (including FOXM1, ETS2, FOXO1). Several of the differential gene targets of these TFs (i.e. *PIK3CD*, *TNFRSF13C, PTPN7, GAB1* and *TCF4)* are conserved in CRLF2 High versus Low patient-derived cell lines and have been previously implicated in leukemias. The network-based approach we applied has been similarly implemented in several other systems to infer regulatory interactions that have been experimentally validated^27–30^. To validate TF-gene inferred interactions, we analyzed CRISPRi, siRNA knockdown, and ChIP-seq ENCODE datasets in K562 and GM12878 cells involving 9 individual TFs. This resulted in the validation of gene targets for each TF further supporting the network-based approach applied in this study. Thus, we uncovered a number of genes along with their regulators that could contribute to the maintenance of leukemia and which act as potential candidates for therapeutic targeting in *CRLF2*-High B-ALL.

## RESULTS

### Genome-wide transcriptional changes in *CRLF2-*overexpressing primary patient samples

Traditional chemotherapy is non-specific and targets rapidly dividing cells. Thus, toxicity results in injury to healthy cells, causing further morbidity^31^. To reduce overall toxicity and improve prognosis, most research has been directed towards understanding the underlying molecular pathology of the leukemia. In this study, we focused on identifying regulatory interactions associated with overexpression of *CRLF2* with the goal of finding new potential therapeutic targets in this subset of B-ALL patients.

Gene expression profiles associated with *CRLF2* overexpression were identified by comparing RNA-sequencing from 15 CRLF2-overexpressing patients (CRLF2-High) and 15 patients with low levels of *CRLF2* (Other B-ALL). Principal component analysis was performed on gene expression levels across all 30 primary patient samples. This revealed a clear separation between the two groups of patients (**Fig 1b**). Principal component analysis with samples labeled according to batch do not demonstrate a batch effect (**Supplementary Fig. 1a**).

To further analyze the changes in gene expression between *CRLF2*-High and other B- ALL patient samples we performed differential expression analysis using DESeq2^32^. This analysis uncovered a total of 3,586 de-regulated genes with an adjusted p-value of less than 0.01 and |log_2_FoldChange| >2 (**Fig. 1c, Supplemental Table 1**). Of these 1470 genes (∼41%) were up-regulated and 2116 (∼59%) down-regulated in the *CRLF2*-High patients. In addition, hierarchical clustering of the 3,586 differentially expressed genes shown in the **Figure 1d** heatmap highlights the prominent transcriptional changes associated with overexpression of *CRLF2*.

*CRLF2*-High patient samples clearly exhibit up-regulation of *CRLF2* (log_2_FoldChange= 10.12) compared to control B-ALL patient samples as shown by RNA-seq tracks on IGV (**Supplementary Fig. 1b**). Based on previous findings we expected to find evidence for activation of JAK-STAT signaling. Indeed, in *CRLF2-*High samples *SOCS2*, a downstream target of JAK/STAT signaling which encodes a SOCS protein that inhibits cytokine signaling as part of a classical feedback loop^33^, was statistically significantly up-regulated (log_2_FoldChange= 4.30) (**Supplemental Table 1**). Thus, consistent with previous findings^34^, JAK-STAT^3, 4^ signaling appears to be activated in these samples.

GSEA was applied on the normalized expression counts of the 3,586 differentially expressed genes using the molecular signatures (MSigDB) database to identify enriched hallmark gene sets that represent well-defined biological processes. Significant pathways with p-value<0.01 and FDR=1% (**Supplemental Table 2**) were enriched including IL2-STAT5 and cytokine mediated signaling (**Supplemental Fig. 1c-1h**).

### *CRLF2*-overexpression leads to promoter-enriched ATAC-seq changes in patient samples

To further investigate the impact of *CRLF2* overexpression, we asked whether CRLF2- High associated transcriptional changes were accompanied by changes in chromatin accessibility. For this we analyzed ATAC-sequencing and compared chromatin accessibility in *CRLF2*-High versus other B-ALL patient samples (7 *CRLF2*-High and 6 other B-ALL). PCA analysis on normalized ATAC-seq signal clearly separated the two conditions (**Figure 2a**). Differential analysis of all the ATAC-seq peaks identified 700 regions that were more accessible and 978 regions with reduced accessibility in the *CRLF2*-High versus low (**Fig. 2b-c**).

**Figure 2:**
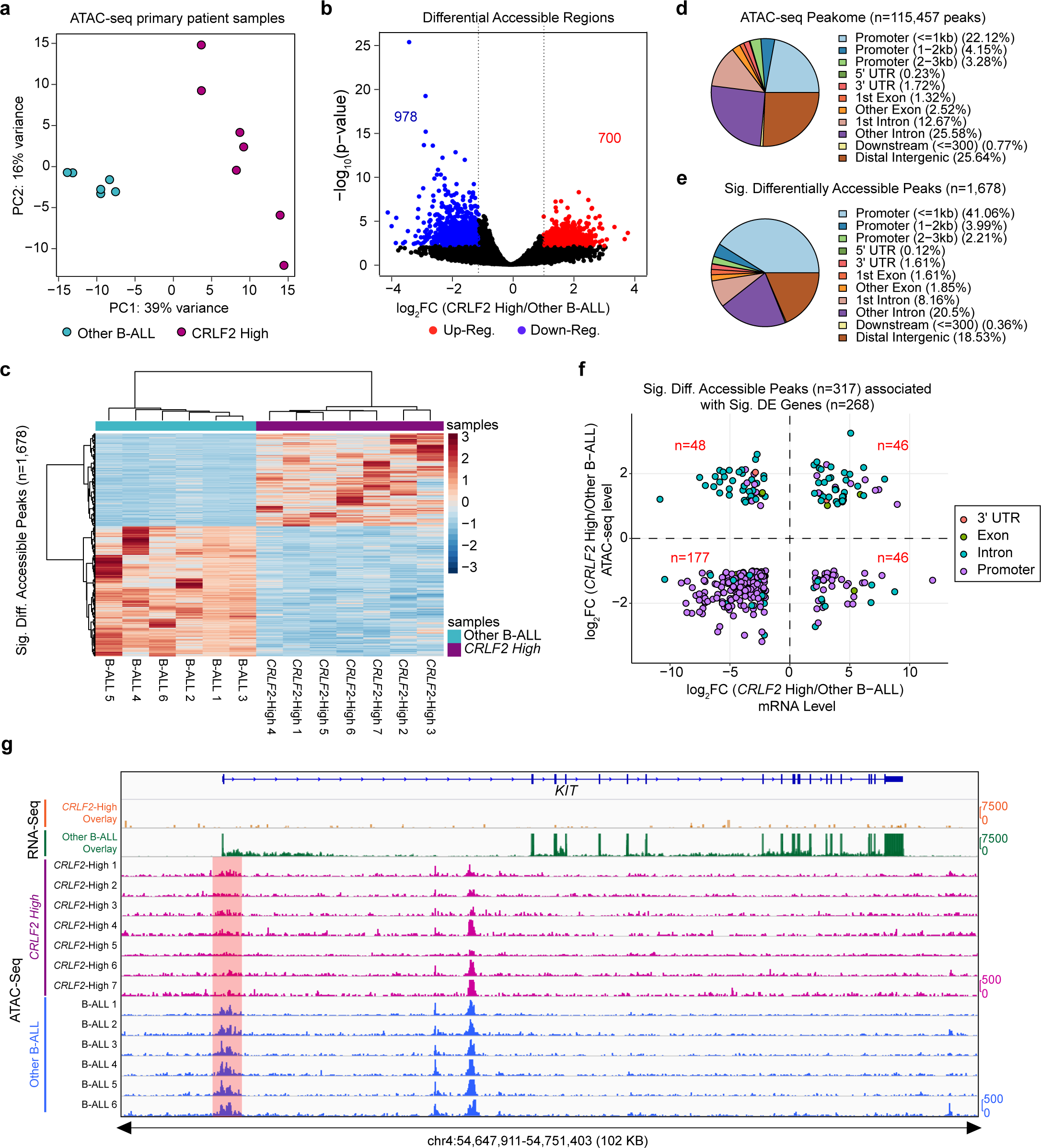
CRLF2-overexpression leads to promoter-enriched ATAC-seq changes in primary patient samples. (a) PCA analysis of 7 CRLF2-High (purple) and 6 Other B- ALL (teal) ATAC-seq samples. (b) Volcano plot of the differential accessible peaks (DESeq2: FDR=1%, |log2 fold change| >1) between CRLF2-High and other B-ALL patients. Red and blue points correspond to more accessible and less accessible ATAC-seq peaks, respectively (n=700 and 978,) in CRLF2-High samples. (c) Heatmap of 700 more accessible (purple bar) and 978 less accessible ATAC-seq peaks in CRLF2-High patients. (d) ChIPseeker genomic annotation of all ATAC-seq peaks or Peakome (n=115,457) across all patient samples. (e) ChIPseeker genomic annotation of significantly differential ATAC-seq peaks (n=1678). (f) Differentially ATAC-seq peaks (n=317), excluding peaks in distal intergenic regions, linked to differentially expressed genes (n=268) and the log2 fold change of gene expression (x-axis) plotted against the log2 fold change of ATAC-seq reads (y-axis). Linked genes are colored according to genomic annotation. (g) IGV screenshot of ATAC-seq tracks for individual samples at down-regulated KIT gene, RNA-seq tracks for CRLF2-High (orange) and other B-ALL (green) samples are overlayed. The highlighted region indicates a region of decreasing accessibility in CRLF2-High samples.

A total number of 115,457 ATAC-seq peaks across all patients were assigned to promoters (within 3kb of TSS), UTRs, exons and introns of genes (**Fig. 2d**). Significant differential ATAC-seq peaks (n=1,678) were similarly annotated (**Fig. 2e**) and more highly enriched (∼47%) at promoter regions compared to all peaks (∼29%). There were 317 (18.9%) differential ATAC-seq peaks associated with significant differentially expressed genes (n=268), either on promoters or gene bodies. The majority of the differential ATAC-seq peaks (223) that overlap differentially expressed genes were concordantly regulated, with the exception of 94 ATAC-gene pairs. Discordant ATAC-gene pairs could be representative of silencers and their gene targets. For example, a decrease in accessibility can impair binding of a silencer to its gene target and therefore lead to an increase in that gene’s expression (**Fig. 2f**). Furthermore, about 79% (177/223) of concordant pairs represented ATAC-seq peaks with decreasing accessibility associated with genes exhibiting reduced expression. An example of this is shown in **Figure 2g**, where an ATAC-seq peak with significantly decreased accessibility is associated with down-regulation of receptor tyrosine kinase *KIT* in *CRLF2*-High patients. Importantly, the majority of transcriptional changes were not associated with significant changes in chromatin accessibility near the promoter or within the gene bodies (3328/3596 genes). In sum, overexpression of *CRLF2* leads to genome-wide chromatin accessibility changes enriched at promoter regions with a small proportion linked to significant transcriptional changes.

### Construction of a B-ALL regulatory network using ATAC-motif derived priors

*CRLF2* overexpression activates a signaling cascade that involves many TF regulators and gene targets. To examine these connections, we wanted to first identify which TFs could potentially be regulating the differentially expressed genes in *CRLF2*-High versus other B-ALL patient cohorts, and second to infer the relationship between these TFs and their target genes.

To address this, we performed TF-motif analysis, using Analysis of Motif Enrichment (AME)^35^ to identify enriched TF motifs at promoter-annotated differentially accessible ATAC-seq peaks compared to shuffled input control sequences. Using an adjusted fisher p-value<1^e-13^, 138 enriched TF motifs were defined at differentially accessible promoter peaks that could be important for the *CRLF2*-High associated gene signature (**Supplemental Fig. 3a**).

**Figure 3:**
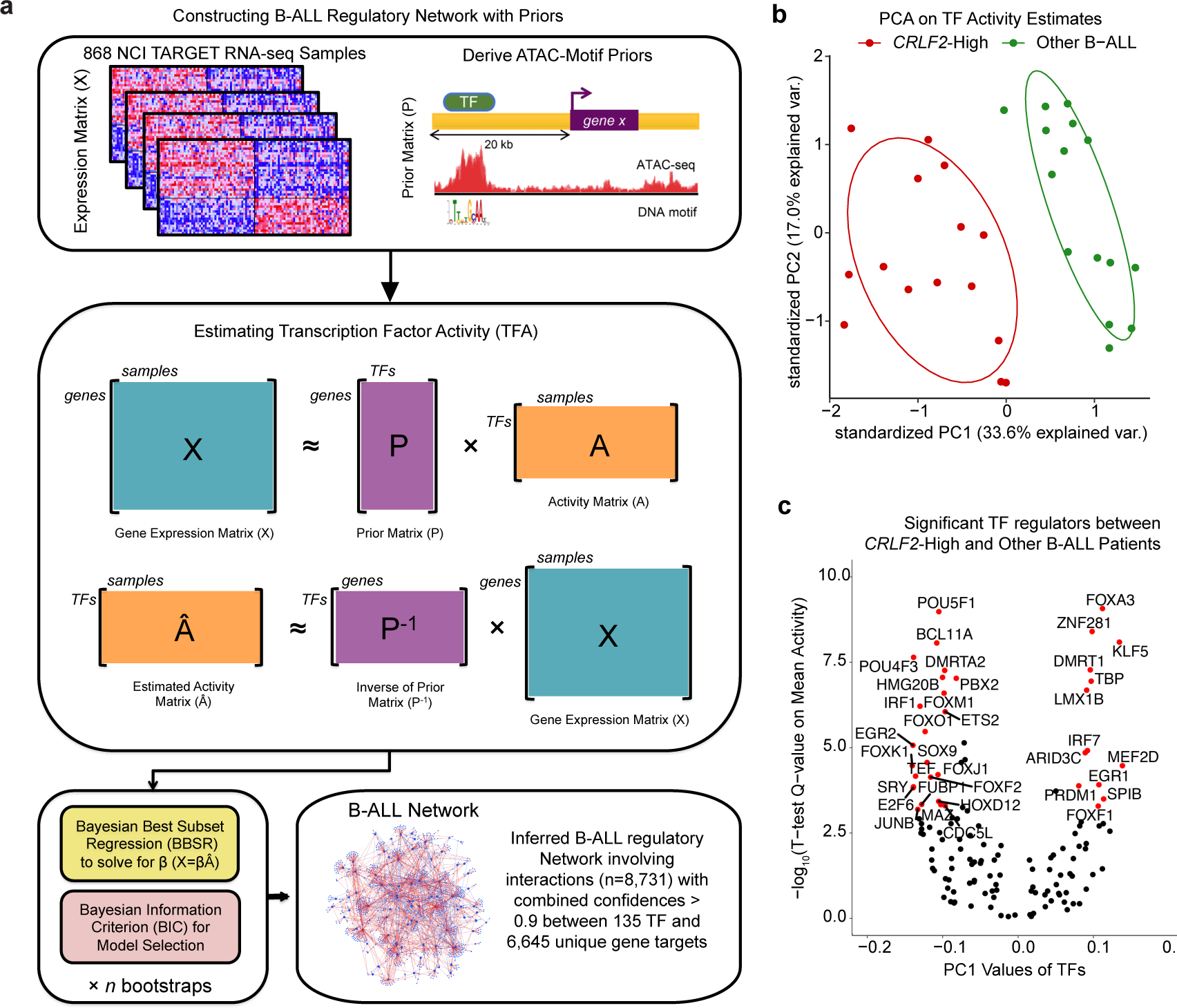
B-ALL network inference identifies significant TFs in the CRLF2-High patient cohort. (a) Workflow for construction of B-ALL regulatory network using Bayesian Best Subset Regression with the Inferelator algorithm. Inferelator resulted in a B-ALL regulatory network involving interactions (n=8,731) with combined confidences > 0.9 between 135 TF and 6,645 unique gene targets. (b) PCA of 30 primary patient samples computed according to estimated TF activity values of 135 TFs. (c) Volcano plot of the significant differential TFs (FDR=0.1%, |PC1 value| >0.075) between CRLF2- High and other B-ALL patients. Red points indicate highly variable TFs between two conditions with significant differential mean activity (t-test; q-value<0.001).

To better understand the relationship between candidate TF regulators identified in **Supplemental Fig. 3a** and the genes they potentially regulate, we sought to infer a transcriptional regulatory network using the Inferelator algorithm^25, 26^. The main limitation of network inference is the sample size (the limited number of samples with an *CRLF2-*High rearrangement). To model transcriptional interactions in a B-ALL specific context requires a large dataset that includes hundreds of B-ALL patient samples, not limited to samples with a specific genetic alteration. For this we made use of the TARGET initiative and analyzed 868 B- ALL RNA-seq samples that were used to obtain a normalized gene expression matrix that could be used for the construction of a B-ALL specific regulatory network.

There are three major steps involved in inferring a transcriptional regulatory network, including generating priors, estimating transcription factor activities, and modeling gene expression with regularized regression. To generate priors, recent studies have incorporated prior information from different data types, like ChIP-seq, ATAC-seq, and TF-motif analysis to associate TFs to their putative gene targets and this has considerably improved network inference ^27–29, 36^. Thus, we focused on combining chromatin accessibility data from the *CRLF2*-High and Low patients with TF-motif analysis to generate priors and acquire the TF-gene associations. First, all ATAC-seq peaks found within a 20kb window upstream of a gene’s transcription start site (TSS) were assigned to that gene. Second, FIMO was used to analyze all known TF motifs enriched within gene-associated ATAC-seq peaks. Lastly, TF motifs were aggregated across all ATAC-seq peaks for each gene. For each TF motif-gene association, a score, a p-value and an adjusted p-value was computed. Only TF-gene associations that had an adjusted p-value less than 0.01 were retained (**Fig. 3a**). With a prior matrix and a normalized gene expression matrix from hundreds of B-ALL specific patient samples, we used the Inferelator algorithm to infer regulatory interactions. Finally, to reduce the number of regulatory hypotheses on TF-gene interactions and improve predictions, we tested only regulatory interactions of TFs (n=138) that were significantly enriched at differentially accessible promoters between *CRLF2*-High and other B-ALL patient samples **(Supplemental Fig 3a)**.

The second step of network inference involves estimating the activity of all TFs in our network based on the cumulative effect of each TF on its target genes^29, 37^. The prior matrix represents a list of potential TF-gene interactions derived from ATAC-TF-motif analysis and the activity of each TF can be estimated by taking the pseudo-inverse of the prior multiplied by the gene expression matrix (**Fig. 3a**). Thus, transcription factor activities were estimated using the ATAC-seq motif derived priors and the gene expression matrix obtained from the TARGET database (**Fig. 3a**). Finally, gene expression was modeled as a weighted sum of its activities using regression with regularization (see methods section for details). The output is a matrix where each row represents a TF-gene interaction, with a β parameter that corresponds to the magnitude and direction of a particular interaction and a computed confidence score (see methods section for details).

Using the ATAC–motif prior, the performance of our model selection within the Inferelator was evaluated by applying a 10-fold cross validation technique. Specifically, we split the prior into 10 equal sized sets. One set was retained as the gold standard to test the model, while the remaining 9 sets were used as the prior. The cross-validation procedure was repeated 10 times, until each of the 10 partitions was used exactly once as the gold standard. As a negative control, we employed the same 10 fold cross-validation strategy after randomly shuffling the prior. We computed the area under the precision-recall curve (AUPR) for each of the 10 iterations and compared the results between the prior and the shuffled prior. Overall, AUPR were significantly higher using the ATAC-seq derived motif prior compared to the randomly shuffled prior (**Supplemental Fig 3b**). Thus, the final network was inferred using the ATAC-seq derived motif prior and the number of significant regulatory interactions was limited to TF-gene interactions with combined confidences > 0.9 (high-confidence interactions). This generated a B-ALL specific regulatory network involving 135 TFs (that may be important regulators of *CRLF2*-High gene signature) and 8,731 high confidence patient specific TF-gene interactions (**Fig. 3a**).

### Significant Transcription Factors altered by *CRLF2* overexpression

To identify TFs affected by *CRLF2* overexpression, we reapplied the TF activity estimation strategy described above to estimate activity across the 30 *CRLF2*-High and other B- ALL samples. By assigning the inferred 8,731 patient-specific TF-gene interactions as priors with the normalized gene expression matrix including all 30 primary patient samples, we estimated an activity matrix of all 135 TFs. Next, we evaluated TFA estimates using PCA analysis across the 30 samples and demonstrated a clear separation of *CRLF2*-High and Low patient samples (**Fig. 3b**). To define the top TFs that explain the variance associated with overexpression of *CRLF2*, we extracted and ranked the absolute value of the PC1 values for each TF (**Supplemental 4a)**. As a secondary means to identify TFs altered at the activity level, we computed the mean activity level of each TF between *CRLF2*-High and Low samples and tested for statistical difference. To narrow the list of potential TFs altered as a result of *CRLF2* overexpression, we ranked the most highly variable TFs and retained those with |PC1| > 0.075, as well as TFs with significant differences in mean activity level with an adjusted p-value 0.001. This resulted in 36 significant TFs as highlighted in the volcano plot in **Fig. 3c**. Boxplots depicting highly variable TFs (n=36) with significant differential mean activities are represented in **Supplemental Fig 4b**. As a result, we defined 36 significant TF regulators predicted to regulate 2,410 gene targets from the inferred B-ALL regulatory network previously described, with the number of targets for each TF shown in **Supplemental Fig 4c**. We note that many TFs (FOXM1^38^, ETS2^39, 40^, BCL11A^41^, FOXO1^42^, etc.) with significantly altered activity are implicated in cancer and could therefore potentially contribute to tumorigenesis in *CRLF2*-High B-ALL patients. Furthermore, 6 of the 36 significant TFs (*ETS2, SRY, FOXO1, EGR1, MEF2D, KLF5*) exhibit a statistically significant difference in mRNA expression between the two patient groups (**Supplemental Fig. 4d**). According to both transcriptional and predicted activity levels, these TFs represent robust candidates that could be important downstream effectors of the *CRLF2*- mediated signaling cascade driving leukemogenesis.

**Figure 4:**
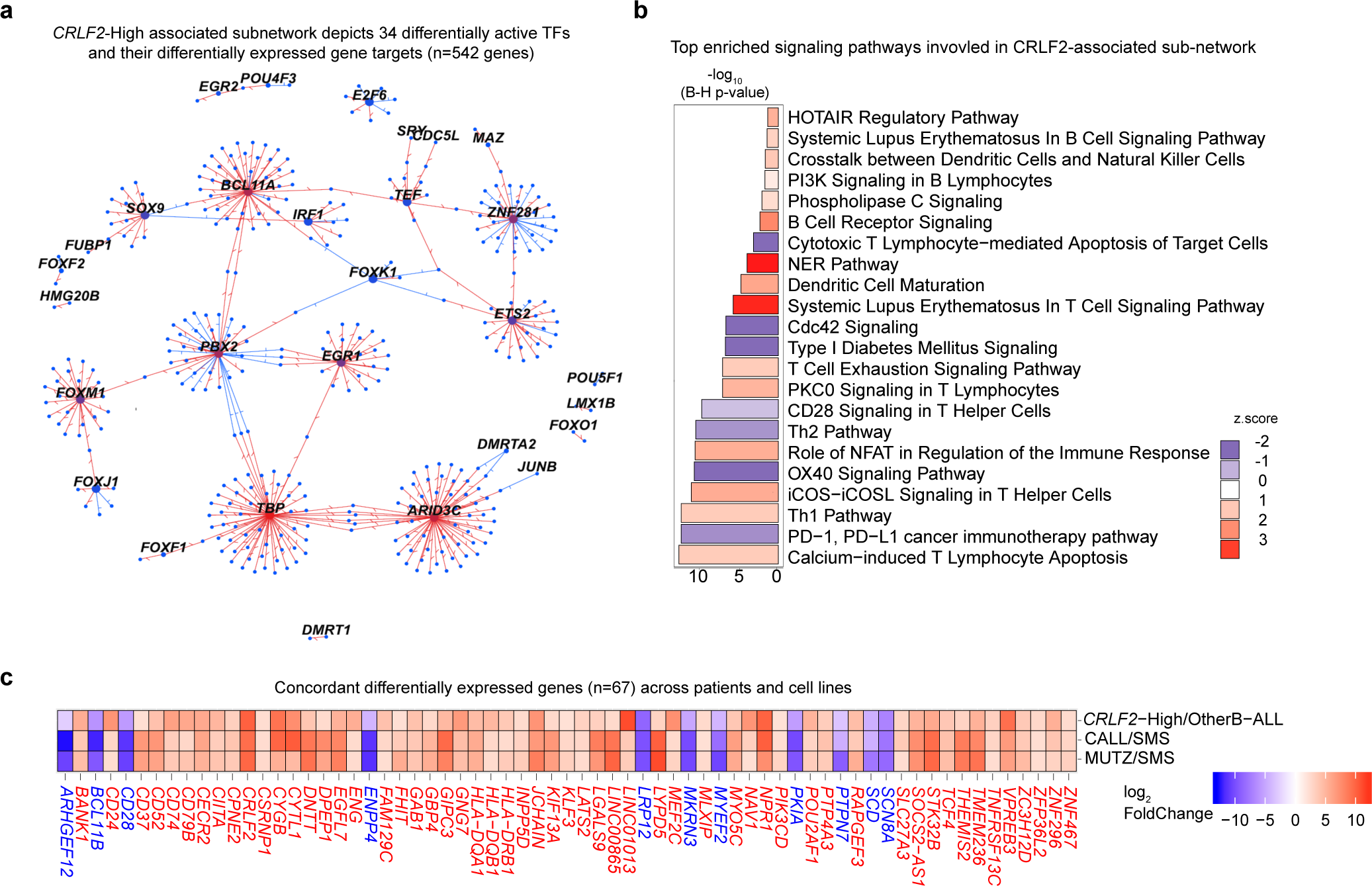
Significant TFs predicted to regulate conserved differentially expressed gene targets across patients and cell lines. (a) Filtered B-ALL network involving regulatory interactions between significant TFs (n=34) and their differentially expressed gene targets (n=542). TFs are the source nodes labeled in black and the gene targets are the blue target nodes. The size of the source nodes reflects the number of targets each TF has. TF activation and repression of a gene are represented by red and blue edges, respectively. (b) Significant pathways (–log10(B-H p-value > 1.3 & |z-score| > 1) identified from analysis of differential gene targets in the CRLF2 specific sub-network are shown at the top of the figure, ranked by the –log10(Benjamini-Hochberg p-value). Z-score in the bar plot indicates sign of the pathway with red and blue representing up- and down-regulation of the pathway. (c) Log2 fold changes of all the convergent differentially expressed gene targets in CRLF2-High vs other B-ALL patients and cell lines (CALL vs SMS, MUTZ vs SMS).

### A *CRLF2-*associated sub-network of altered TFs and their differentially expressed targets

To filter regulatory interactions to a specific *CRLF2*-High sub-network, we computed the number of gene targets that were differentially expressed (542/2410) in *CRLF2*-High versus *CRLF2*-low patient samples (**Fig. 4a**). Ingenuity pathway analysis was performed on the 542 differentially expressed gene targets in the *CRLF2*-High associated sub-network. This generated 22 significant pathways (**Fig. 4b**) including the activation of the canonical PI3K pathway (**Fig. 4a**). The majority of the gene targets are involved in more than one signaling pathway and many of these pathways are important in adaptive immunity and inflammatory responses.

To define a set of genes in the *CRLF2*-High associated sub-network that could have the highest potential as therapeutic targets, we turned to patient-derived cell lines to identify robust changes across both model systems. The patient-derived cell lines used in our analysis were MHH-CALL-4^43^ (CALL) and MUTZ5^44^ (MUTZ) that harbor *IGH-CRLF2* translocations leading to *CRLF2* overexpression, and a non-translocated B-ALL pre-B leukemic cell line, SMS-SB^45^ (SMS). As seen in the RNA-seq tracks of **Supplementary Fig. 5a**, *CRLF2* is clearly overexpressed in the translocated CALL and MUTZ cell lines, compared to non-translocated SMS. Principal component analysis of cell line RNA-seq samples (**Supplementary Fig. 5b**) indicated that the majority of the variance (71%) lies between *CRLF2*-High cell lines and SMS cell lines, while about 26% of variance separated the two *CRLF2*-High cell lines. Thus the PCA analysis demonstrates overexpression of *CRLF2* is the major difference between the two conditions, consistent with what was observed in patient samples.

Differential gene expression analysis of CALL versus SMS (1,631 DE genes) and MUTZ versus SMS (1,484 DE genes) identified thousands of differentially expressed genes (**Supplementary Fig. 5c**), with roughly equivalent numbers of up and down-regulated genes in each case. The number of overlapping differentially expressed genes is shown in **Supplementary Fig. 5d**. As shown in **Supplementary Fig. 5e,** the log_2_ fold change of (CALL/SMS) and (MUTZ/SMS) demonstrated that about 97% of the genes were in convergent orientation (561 up-regulated, 353 down-regulated). We compared the gene targets in the *CRLF2*-associated regulatory sub-network with the convergent differentially expressed genes in cell lines and identified 67 significant convergent transcriptional alterations that were conserved across *CRLF2*-High patient and cell line samples. This is shown in **Figure 4c** by the log_2_ fold change of all 67 convergent differentially expressed genes in the patients and each pairwise cell line comparison (CALL versus SMS, and MUTZ versus SMS). Of the 67 gene targets, 84% are up-regulated in *CRLF2*-High samples including *PIK3CD, DPEP1, PTPN7, TNFRSF13C (BAFFR)* and Class II major histocompatibility complex (MHC) genes. Previous studies have implicated the majority of these genes in hematologic malignancies including B-ALL^46–51^ but their affect specifically in *CRLF2-*overexpressing B-ALL is not known. Furthermore, thirteen of these genes have available inhibitors (**Supplemental Table 3**).

### Validating Inferred Gene Targets of TFs significantly altered by *CRLF2* overexpression

To support our network-based approach in identifying TF-gene regulatory interactions in leukemic cells, we sought to validate our predictions of interactions involving TFs using publicly available data in similarly related cells. We collected ENCODE data to support TF-gene associations involving 9 significant TFs. The experiments analyzed include CRISPR interference (CRISPRi) targeting of FOXM1 in myelogenous leukemic cells (K562), siRNA targeting of MAZ and E2F6 in K562 cells, and ChIP-seq for 6 significant TFs (BCL11A, ETS2, EGR1, PBX2, IRF1, TBP) in K562 cells and a B-lymphocyte derived cell line (GM12878).

First, we computed the number of gene targets we inferred for each of the 9 TFs (**Figure 5a**). To validate some of the predicted gene targets of FOXM1, we compared RNA-seq samples in K562 cells treated with CRISPRi targeting of FOXM1 versus non-targeted cells. A PCA of these samples distinctly separates the FOXM1 CRISPR targeted samples from the non-targeting control samples (**Figure 5b**). DESeq2 was applied was used to examine differentially expressed genes associated with FOXM1 targeting. These were overlapped with the network’s predicted gene targets of FOXM1. Of the 123 gene targets we predicted to be regulated by FOXM1, 50 (40%) had a significant change in their expression upon FOXM1 CRISPRi targeting including *CRLF2* itself (**Figure 5c**). FOXM1 is a forkhead box transcription factor functioning downstream of the PI3K/AKT/mTOR pathway, which is known to be induced in *CRLF2*-overexpressing B-ALL^7, 52^. It is involved in cell proliferation, DNA repair, and self-renewal.

**Figure 5:**
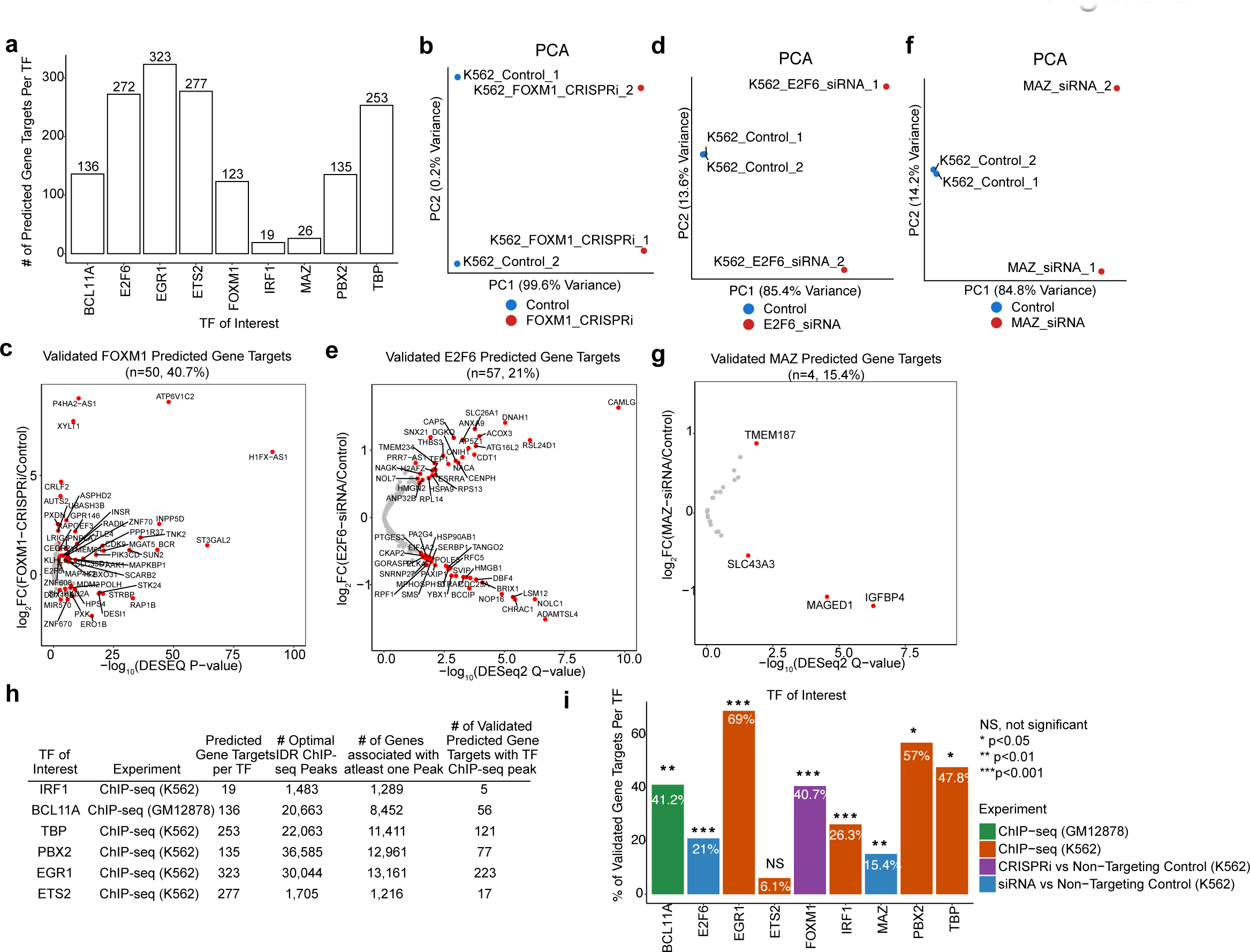
Validating inferred gene targets of TFs significantly altered by *CRLF2* overexpression. (a) Bar plot representing the number of predicted gene targets for 9 TFs, for which public data was available for validation. Exact numbers are represented at the top of the bar. (b) PCA analysis of 4 CRISPRi targeting FOXM1 (red) and non-targeting control (blue) RNA-seq samples. (c) Volcano plot of the FOXM1 predicted gene targets, those that are differentially expressed genes (FDR=5%, |log2 fold change| >0.5) between FOXM1 CRISPRi and non-targeting control samples are highlighted in red and labeled. (d) PCA analysis of 4 CRISPRi targeting E2F6 (red) and non-targeting control (blue) RNA-seq samples. (e) Volcano plot of the E2F6 predicted gene targets, those that are differentially expressed genes (FDR=5%, |log2 fold change| >0.5) between E2F6 CRISPRi and non-targeting control samples are highlighted in red and labeled. (f) PCA analysis of 4 CRISPRi targeting MAZ (red) and non-targeting control (blue) RNA-seq samples. (g) Volcano plot of the MAZ predicted gene targets, those that are differentially expressed genes (FDR=5%, |log2 fold change| >0.5) between MAZ CRISPRi and non-targeting control samples are highlighted in red and labeled. (h) Table representing the ENCODE ChIP-seq experiments for 6 TFs, including the TF of interest, the experiment, number of predicted gene targets from the B-ALL network, number of optimal IDR ChIP-seq peaks, number of annotated genes with at least one ChIP-seq peak, and finally the number of validated gene targets for each TF of interest. (i) Barplot representing the percentage of validated gene targets for each of the 9 TFs of interest. The percentage of validated inferred gene targets for each TF is labeled in white at the top of the bar. P-values obtained from hyper-geometric test are annotated as follows: NS, not significant; *, p < 0.05, p<0.01, p<0.001. Color of the bar represents the experiment.

Overexpression of FOXM1 has been detected not only in B-ALL^38^ but also in a wide range of human cancers and its overexpression is linked to poor prognosis^38, 53–62^.

To evaluate predicted gene targets of another significant TF, we identified genes altered by siRNA knockdown of the E2F6 gene in K562 cells. E2F6 is one of the E2F transcription factors that play distinct and overlapping roles in regulating cell cycle progression^63^. E2F proteins can be transcriptional activators or repressors. In particular, E2F6, has been shown to repress the transcription of known E2F target genes and is considered a dominant negative inhibitor of the other E2F proteins. Yeast two-hybrid assays have determined that E2F6 associates with known components of the polycomb complex, suggesting a role for E2F6 in recruiting the polycomb transcriptional repressor to target genes^64^. To support some the inferred gene targets of E2F6 using the network based strategy in this study, we analyzed E2F6 targeted RNA-seq samples using siRNA versus non-targeted controls. A PCA analysis confirmed expected clustering of replicates for E2F6 targeted and control samples (**Figure 5d**). Differentially expressed genes were determined and overlapped with inferred gene targets of E2F6. This analysis supported 57 of the 272 (21%) predicted E2F6 gene targets (**Figure 5e**).

Next, we analyzed predicted gene targets of the MAZ TF, also known as Myc-associated zinc-finger protein. This TF is ubitiquously expressed in most tissues and has been shown to activate a variety of genes including c-Myc, insulin, VEGF, and MYB, and repress the p53 protein^65–67^. Specifically, it has been shown that the kinase Akt phosphorylates MAZ, which leads to the release of the MAZ TF from the p53 promoter followed by p53 transcriptional activation^66^. MAZ has also recently been identified as a factor that interacts with cohesin and through this connection alter gene regulation as a result of its impact on TAD structure^68^. As MAZ has been shown to have a dual role in both activation and repression of cancer-related genes, it’s role in different cancer types can vary greatly depending on its target genes. In this particular study, we have predicted significant B-ALL specific regulatory interactions between MAZ and 26 gene targets. To provide evidence supporting these predictions, we analyzed siRNA knockdown of MAZ in K562 RNA-seq samples compared to control samples. A PCA on MAZ knockdown and control K562 samples indicated reproducible replicates that cluster according to their condition (**Figure 5f).** Genes that were differentially expressed were analyzed and overlapped with predicted MAZ gene targets, resulting in 15% of predicted gene targets’ expression significantly altered by siRNA knockdown of MAZ (**Figure 5g).**

Finally, to support TF-gene regulatory interactions of other significant factors, we downloaded available ENCODE optimal IDR (Irreproducible Discovery Rate) peaks for 6 important TFs (BCL11A, ETS2, EGR1, PBX2, IRF1, TBP) in K562 and GM12878 cell lines. Optimal IDR peaks represent the most highly reproducible peaks across replicates. The aim here was to delineate potential gene targets of each TF by determining binding sites according to significant and reproducible ChIP-seq peaks at promoters. Thus, ChIP-seeker was applied to annotate the optimal IDR peak sets of each TF to genes, using the same threshold applied to generate TF-gene priors as previously described (20 KB window upstream of the TSS), the number of genes associated with at least one ChIP-seq peak for each TF is shown in **Figure 5h**. For each individual TF of interest, we overlapped the network inferred gene targets of that TF with the genes associated with ChIP-seq binding. Overall, we were able to validate 40.1% of BCL11A gene targets, 69% of EGR1 gene targets, 6.1% of ETS2 gene targets, 26.3% of IRF1 gene targets, 57% PBX2 gene targets, and 47.8% of TBP gene targets that we infer using the network-based approach (**Figure 5i**). Finally, individual hyper-geometric tests were performed for each experiment; to test the significance of the proportion of validated gene targets compared to random. The proportion of validated gene targets was significant for almost all 9 TFs, with the exception of ETS2, which could most likely be a matter of cell-type context (**Figure 5i**).

## DISCUSSION

Patients with B-ALL who overexpress *CRLF2* are at increased risk for refractory disease and relapse. This alteration leads to the activation of JAK-STAT and other associated pathways^3^. Available treatments include non-specific cytotoxic chemotherapies which though often effective are responsible for many of the toxicities seen in these patients. The addition of Tyrosine-Kinase Inhibitors and JAK inhibitors in the treatment of leukemias has allowed for targeted treatments but these treatments also affect the biology of healthy cells. Furthermore, the use of *CRLF2*-targeted CAR T cells shows great promise in the eradication of the cancer, yet limitations and toxicities associated with this treatment have yet to be studied in patients. Thus, it is important to more deeply investigate the regulatory control underlying *CRLF2*- overexpressing leukemia to identify and evaluate alternate gene therapeutic targets for tailored treatment.

In this work, we sought to better understand the impact of *CRLF2* overexpression on leukemogenesis, with the aim of delineating the regulatory interactions for patient-specific *CRLF2-*High B-ALL. We analyzed RNA-seq from *CRLF2*-High and Other B-ALL patient samples and identified hundreds of differentially expressed genes. We subsequently analyzed ATAC-seq data to evaluate the effects of this genetic alteration on DNA accessibility. We found accessibility to be altered across the genome with an enrichment of changes localized to promoter regions. Only a small proportion of these altered promoter regions are linked with gene expression changes.

To systematically annotate regulatory interactions, we inferred putative binding of specific TF motifs at all ATAC-seq peaks found at the promoters of genes in the genome.

Leveraging hundreds of available B-ALL RNA-seq samples from the TARGET database along with prior knowledge of TF-motifs at accessible promoters allowed us to infer a B-ALL specific regulatory network involving TF motifs (135) enriched at differentially accessible promoters. Deeper investigation of the TF regulators and robust differential gene targets specific to the *CRLF2*-High sub-network identified a candidate list of potential therapeutic targets.

*PIK3CD*, one of the significantly overexpressed candidates identified in *CRLF2-*High patients, encodes a class I phosphoinositide-3 kinase (PI3Ks), which is highly enriched in leukocytes and known as p110δ. Up-regulation of this gene has been demonstrated to occur in T-cell ^46^ acute lymphoblastic leukemia and it has emerged as a therapeutic target. Specifically, the use of a p110δ inhibitor has led to reduced proliferation and apoptosis in T-ALL cell lines^46^. Furthermore, this gene target has been similarly implicated in Philadelphia chromosome–like acute lymphoblastic leukemia (Ph-like ALL), another high-risk B-ALL subtype correlated with poor prognosis^14^. *CRLF2* overexpression occurs in about 50% of Ph-like ALL cases. Using patient-derived xenograft (PDX) models of childhood Ph-like ALL, Tasian et al. demonstrate the therapeutic efficacy of Idelalisib, a PI3Kδ inhibitor in vivo^14^. That PIK3CD has been previously implicated in *CRLF2*-overexpressing B-ALL supports our approach in identifying relevant *CRLF2*-associated gene targets.

Another gene target up-regulated in *CRLF2*-High patients is *GAB1*. This gene encodes an adapter protein that recruits proteins, including PI3K to the plasma membrane to propagate signals for many important biological processes such as cell proliferation, motility, and erythrocyte development. *GAB1* proteins lack enzymatic activity, but are phosphorylated at tyrosine residues via interaction with PI3K, and this association is essential for activation of the PI3K/AKT and MAPK/ERK pathways^69^. Up-regulation of *GAB1* has been linked to breast cancer progression^48^ and implicated in BCR-ABL positive ALL cells^47^. Previous analysis of B-ALL adult patients with the BCR-ABL alteration versus BCR-ABL–negative adult ALL patients, not only identified overexpression of *GAB1* but also up-regulation of several class II major histocompatibility complex (MHC) genes and their trans activator *CIITA*^47^. These results are similarly reproduced in our own data demonstrating conserved co-expression of *GAB1*, *CIITA, HLA-DQA1, HLA-DQB1,* and *HLA-DRB1* across patients and cell lines. In addition, we determined that 4 out of 5 of these genes (*GAB1*, *CIITA, HLA-DQA1, HLA-DQB1)* are predicted to be gene targets of the BCL11A transcription factor. BCL11A is a zinc finger protein that is expressed in hematopoietic cells and in the brain and whose presence is linked to hematological malignancies^41, 70^. This TF has been shown to play an important role in lymphoid development. BCL11A expression in adult mouse is found in most hematopoietic cells but enriched in B cells, early T cell progenitors, common lymphoid progenitors (CLPs) and hematopoietic stem cells (HSCs). The deletion of BCL11A leads to early B-cell and CLP apoptosis and impairs the development of HSCs into B, T, and natural killer (NK) cells^71^. BCL11A is a key regulator of fetal-to-adult hemoglobin switching. This regulator is also known to interact with the chromatin remodeling SWI/SNF complex and the nucleosome remodeling and deacetylase (NuRD) complex.^72^. The analysis described earlier, associating BCL11A ChIP-seq binding sites to genes in GM12878 cells, validated 56 of BCL11A’s predicted gene targets, including overexpressed *GAB1*, *CIITA, HLA-DQA1, HLA-DQB1,* and *HLA-DRB1* in both patients and cell lines. Another candidate gene target of BCL11A, validated by ChIP-seq binding, is *DPEP1.* As a zinc-dependent peptidase, *DPEP1* is important for metabolism of the antioxidant glutathione and was recently linked to relapse in B-ALL. Specifically, high levels of this gene were linked with a higher incidence of relapse and worse overall survival^73^.

*GAB1* is also one of the few gene targets that we predict to be regulated by two significantly altered TFs, BCL11A and PBX2. The pre-B-cell leukemia transcription factor 2, PBX2 is a ubiquitously expressed protein and can regulate gene expression and effect processes such as cell proliferation and differentiation^74^. It has also been shown to work as a cofactor with the HOXA9 protein^75^. Knockdown of PBX2 leads to increased apoptosis in adenocarcinoma and esophageal squamous cell carcinoma cancer cell lines^76^. PBX2 is also an inferred regulator of the down-regulated *PTPN7* gene in *CRLF2*-overexpressing patients and cell lines. This gene encodes a protein tyrosine phosphatase that is mainly expressed in hematopoietic cells and known to negatively regulate the activation of ERK and MAPK kinases^77^. Overexpression of *PTPN7* in T-cells down-regulates ERK signaling and reduces T-cell activation and proliferation^78^. Thus, *PTPN7* represents another strong candidate gene target as it can act as a negative regulator of one the canonical signaling pathways downstream of *CRLF2*.

The up-regulation of *TNFRSF13C*, has also been linked to relapse in B-cell acute lymphoblastic leukemia. *TNFRSF13C* encodes the receptor for B-cell activating factor (BAFFR) and is important for B-cell maturation and survival. Expression of *TNFRSF13C* is primarily detected in primary B-cell malignancies and B-cell related organs and absent in other healthy related tissues^79, 80^. There is evidence suggesting the expression of this receptor plays a role in relapse as well as persistence of leukemic cells after early treatment^80^. Current drugs available to target the BAFFR receptor include the Anti–BAFF-R antibody VAY-736^81^. In addition, a CAR-based strategy has recently been engineered to target this receptor. Thus, *TNFRSF13C* represents a potential target for *CRLF2*-overexpressing leukemias.

The network-driven approach also identified a number of significant TF regulator motifs including ETS2^40^, FOXO1^42^, EGR1^82^, and FOXM1^38, 55, 56^, which are implicated in hematological malignancies. FOXM1 could be an important regulator in *CRLF2*-overexpressing cells due to its crucial role in cell proliferation and its implication in various cancer types, as described earlier. Furthermore, our investigations predict that FOXM1 regulates *CRLF2* itself. This is interesting, as FOXM1 is a known regulator functioning downstream of *CRLF2*-associated PI3K/AKT pathway, yet it was not known that FOXM1 could regulate *CRLF2*.

Another candidate regulator, ETS2, a member of the *Ets* family of transcription factors, is involved in cell proliferation and differentiation, and is a known target of the RAS/MAPK signaling pathway^83^. In this study, ETS2 is differentially expressed and predicted to regulate the aforementioned gene target *TNFRSF13C* or BAFFR. In AML, overexpression of ETS2 has been shown to be associated with shorter overall survival and be a predictor of poor prognosis^39^. Analysis of overexpressed ETS2 in AML is associated with co-expression of *SOCS2* and *TCF4*, both of which are significantly up-regulated in *CRLF2*-High B-ALL patient samples. *A*s mentioned previously, *SOCS2* is induced upon cytokine signaling. *TCF4*, also known as *TCF7L2*, is a transcription factor that plays an important role in the Wnt signaling pathway and its overexpression is required for proliferation and survival of AML cells^84^. Indeed, the results of our present study identified *TCF4* as part of the *CRLF2*-associated sub-network and it is overexpressed in both primary patients and patient derived cell lines. Given the role of *TCF4* in the Wnt signaling pathway and its link to poor clinical outcome in AML, this gene represents another putative gene target in *CRLF2*-overexpressing B-ALL.

FOXO1, another significantly altered TF, is one of the FOXO transcription factors regulated by the PI3K/AKT signaling pathway. FOXO TFs are mostly considered as tumor suppressors given their ability to inhibit cancer cell growth. Yet there is contrasting evidence for their activation supporting tumor progression and inducing drug resistance^85^. In particular, the TF FOXO1 plays a vital role in regulating proliferation and cell survival during B-cell differentiation and it’s activity, has been shown to be essential in B-cell precursor acute lymphoblastic leukemia^42^. In this specific study in B-ALL, Wang et al. inhibited FOXO1 and demonstrate reduced leukemic activity in cell lines, primary patients, and xenograft models^42^.

The Early Growth Response 1 (EGR1) TF represents another candidate TF that is not only significantly altered at the gene expression level but also predicted to have a change in activity. This TF is induced in response to B cell receptor (BCR) signaling and mediated by the Ras/Raf-1/MAPK pathway. Studies have also shown that EGR1 can promote differentiation in pre-B cells^86^. High levels of EGR1 have been linked to human diffuse large B lymphoma cells suggesting a role for EGR1 in B lymphoma cell growth^82^. In addition to previous evidence supporting a role for FOXM1, ETS2, BCL11A, PBX2, EGR1, and FOXO1 in other malignancies, the results of our analysis now implicate these TFs in *CRLF2*-overexpressing leukemia. We speculate that downstream of *CRLF2*-mediated signaling, there are alterations to the accessibility landscape that may be influencing TF binding and thereby deregulating transcriptional programs. We have identified a set of TFs and their respect gene targets that we propose may be important factors to consider in *CRLF2*-overexpressing leukemia (**Fig. 6a**). Finally, using CRISPRi and siRNA RNA-seq data for FOXM1, E2F6, and MAZ, as well as ChIP-seq for 6 other TFs from ENCODE, it was possible to validate a proportion of our predictions linking these TFs and their predicted gene targets.

**Figure 6:**
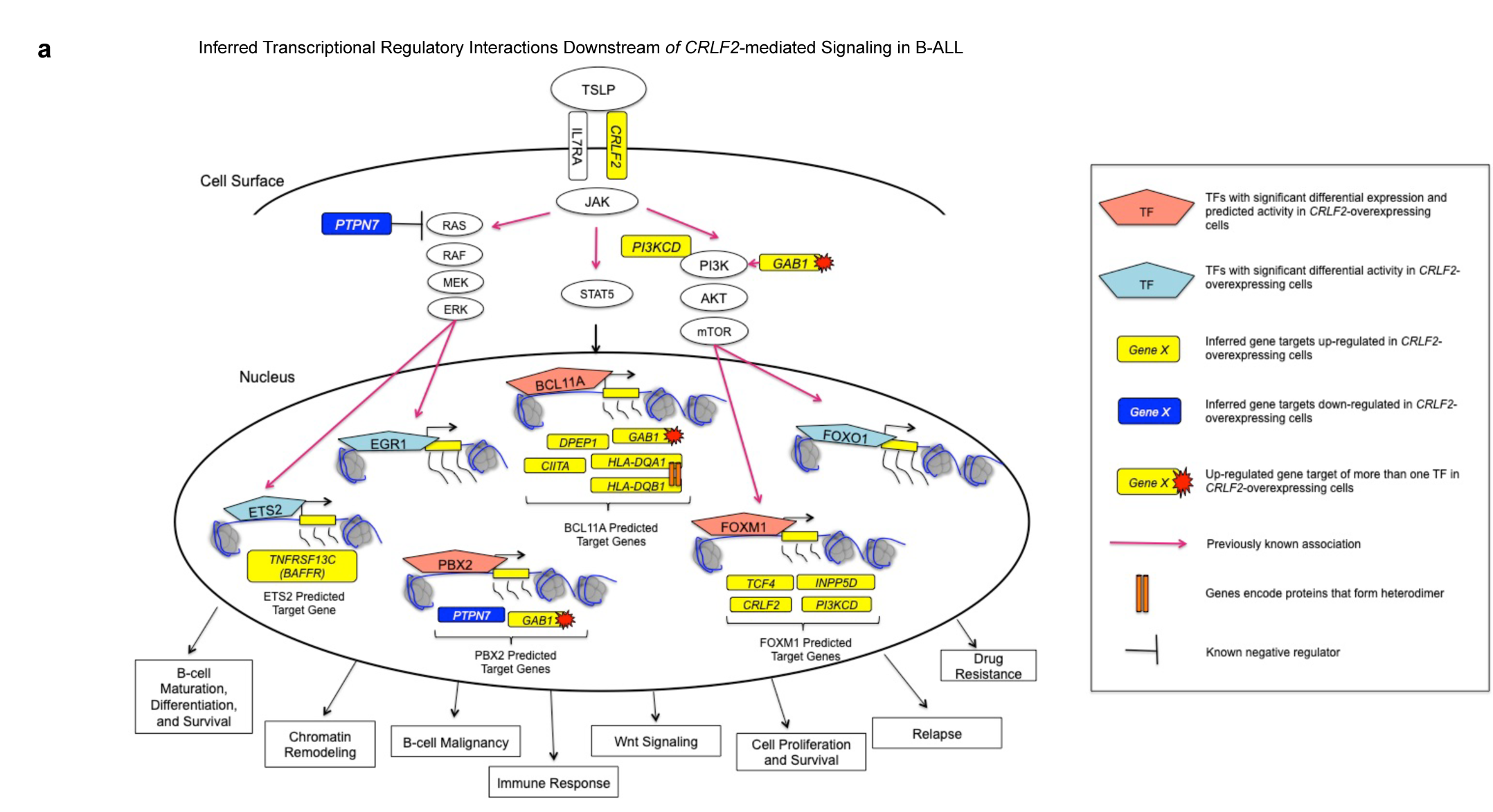
Proposed Model. (a) Inferred transcriptional regulatory interactions downstream of CRLF2-mediated signaling in B-ALL.

In summary, we have investigated the transcriptional impact of *CRLF2* overexpression in a high-risk subset of B-ALL patients. Using an integrative approach, we derived regulatory interactions that have enabled the identification of potential candidates for targeted therapies in B-ALL patients with *CRLF2* overexpression. *PIK3CD, GAB1, DPEP1, TNFRSF13C, TFC4, PTPN7, BCL11A, FOXM1, ETS2, EGR1, PBX2,* and *FOXO1* are amongst the most interesting candidates as there is evidence supporting their role in hematological malignancies. The strategy we have used here can further be adapted to elucidate important regulators and gene targets responsible for additional subsets of B-ALL and other challenging malignancies.

## METHODS

### Cell Culture

The MUTZ5 (RRID:CVCL_1873) and MHH-CALL-4 (RRID:CVCL_1410) human B-cell leukemia cell lines carrying CRLF2-High translocation were obtained, as well as the SMS-SB (RRID:CVCL_AQ30) human B-cell leukemia control cell line. All cells were maintained in RPMI1640 supplemented with 20% FBS and 1% penicillin-streptomycin under 5% CO2 at 37 degrees. Ficoll-enriched, cryopreserved bone marrow samples were obtained from the Children’s Oncology Group (COG) from pediatric patients with B-precursor ALL from biology protocols AALL03B1 and AALL05B1. Both *CRLF2*-HIGH and control patient samples were used. All subjects (or their parents) provided written consent for banking and future research use of these specimens in accordance with the regulations of the institutional review boards of all participating institutions.

### RNA-seq

RNA was extracted using the RNeasy plus kit from QIAGEN. For the cell lines, ribosomal transcripts were depleted using the Illumina® Ribo Zero Module and for the patients, poly- adenylated transcripts were positively selected using the NEBNext® Poly(A) mRNA Magnetic Isolation Module, following the respective kit procedure. Libraries were prepared according to the directional RNA-seq dUTP method adapted from http://waspsystem.einstein.yu.edu/wiki/index.php/Protocol:directional_WholeTranscript_seq that preserves information about transcriptional direction. Sequencing was performed with Illumina Hi-Seq 2500 using 50 cycles paired-end mode.

### ATAC-seq

Cell were counted to collect 50,000 cells per sample. The assay was performed as described previously^87^. Cells were washed in cold PBS and resuspended in 50 µl of cold lysis buffer (10 mM Tris-HCl, pH 7.4, 10 mM NaCl, 3mM MgCl2, 0.1% IGEPAL CA-630). The tagmentation reaction was performed in 25 µl of TD buffer (Illumina Cat #FC-121-1030), 2.5 µl Nextera Tn5 Transposase, and 22.5 µl of Nuclease Free H2O at 37oC for 30 min. DNA was purified on a column with the Qiagen Mini Elute kit, eluted in 10 µl H2O. Purified DNA (10 µl) was combined with 10 µl of H2O, 2.5 µl of each primer at 25 mM and 25 µl of NEB Next PCR master mix. DNA was amplified for 5 cycles and a monitored quantitative PCR was performed to determine the number of extra cycles needed as per the original ATAC-seq protocol^87^. DNA was purified on a column with the Qiagen Mini Elute kit. Samples were quantified using Tapestation bioanalyzer (Agilent) and the KAPA Library Quantification Kit and sequenced on the Illumina Hi-Seq 2000 using 50 cycles paired-end mode.

## QUANTIFICATION AND STATISTICAL ANALYSIS

### RNA-seq analysis in *CRLF2*-High and other B-ALL patient cohort

RNA-seq with replicates for individual patient samples were pooled. Paired-end reads were mapped to the hg38 genome using TopHat2^88^ (parameters:–no-coverage-search–no-discordant–no-mixed–b2-very-sensitive–N 1). To Counts for Refseq genes were obtained using htseq-counts^89^. PCA was performed using R to visualize the clustering of the patient samples.

Differential analysis was performed using DESeq2^32^ and differential genes were considered statistically significant if they had an adjusted p-value less than 0.01 and |log2 FoldChange| >2. Hierarchical clustering was performed on significant differentially expressed genes (3,586) with the manhattan distance and ward.d2 clustering method, using heatmap3 R package for visualization. Bigwigs were obtained for visualization on individual as well as merged bam files using Deeptools^90^ (parameters bamCoverage–binSize 1–normalizeUsing RPKM). Gene Set Enrichment Analysis (GSEA) for pathway enrichment was performed using the default parameters of the GSEA desktop application on normalized reads counts for all patient samples for the 3,586 differentially expressed genes between *CRLF2*-High and other B-ALL patients. See below for information for each RNA-seq sample:

| Original Label | Source | File Name | Sample Name |
| --- | --- | --- | --- |
| CRLF2-High_1 | Skok Lab | CRLF2-High_1_RNA_counts.txt | CRLF2-High_1 |
| CRLF2-High_2 | Skok Lab | CRLF2-High_2_RNA_counts.txt | CRLF2-High_2 |
| CRLF2-High_3 | Skok Lab | CRLF2-High_3_RNA_counts.txt | CRLF2-High_3 |
| CRLF2-High_4 | Skok Lab | CRLF2-High_4_RNA_counts.txt | CRLF2-High_4 |
| CRLF2-High_5 | Skok Lab | CRLF2-High_5_RNA_counts.txt | CRLF2-High_5 |
| CRLF2-High_6 | Skok Lab | CRLF2-High_6_RNA_counts.txt | CRLF2-High_6 |
| CRLF2-High_7 | Skok Lab | CRLF2-High_7_RNA_counts.txt | CRLF2-High_7 |
| CRLF2-High_8 | Skok Lab | CRLF2-High_8_RNA_counts.txt | CRLF2-High_8 |
| CRLF2-High_9 | Skok Lab | CRLF2-High_9_RNA_counts.txt | CRLF2-High_9 |
| CRLF2-High_10 | Skok Lab | CRLF2-High_10_RNA_counts.txt | CRLF2-High_10 |
| CRLF2-High_11 | Skok Lab | CRLF2-High_11_RNA_counts.txt | CRLF2-High_11 |
| CRLF2-High_12 | Skok Lab | CRLF2-High_12_RNA_counts.txt | CRLF2-High_12 |
| SRR457636 | TARGET | CRLF2-High_13_RNA_counts.txt | CRLF2-High_13 |
| SRR457642 | TARGET | CRLF2-High_14_RNA_counts.txt | CRLF2-High_14 |
| SRR457653 | TARGET | CRLF2-High_15_RNA_counts.txt | CRLF2-High_15 |
| SRR3162137 | TARGET | BALL_1_RNA_counts.txt | BALL_1 |
| SRR3162141 | TARGET | BALL_2_RNA_counts.txt | BALL_2 |
| SRR3162143 | TARGET | BALL_3_RNA_counts.txt | BALL_3 |
| SRR3162144 | TARGET | BALL_4_RNA_counts.txt | BALL_4 |
| SRR3162148 | TARGET | BALL_5_RNA_counts.txt | BALL_5 |
| SRR3162151 | TARGET | BALL_6_RNA_counts.txt | BALL_6 |
| SRR3162166 | TARGET | BALL_7_RNA_counts.txt | BALL_7 |
| SRR3162280 | TARGET | BALL_8_RNA_counts.txt | BALL_8 |
| SRR3162282 | TARGET | BALL_9_RNA_counts.txt | BALL_9 |
| SRR3162293 | TARGET | BALL_10_RNA_counts.txt | BALL_10 |
| SRR3162301 | TARGET | BALL_11_RNA_counts.txt | BALL_11 |
| SRR3162304 | TARGET | BALL_12_RNA_counts.txt | BALL_12 |
| SRR3162313 | TARGET | BALL_13_RNA_counts.txt | BALL_13 |
| SRR3162316 | TARGET | BALL_14_RNA_counts.txt | BALL_14 |
| SRR3162374 | TARGET | BALL_15_RNA_counts.txt | BALL_15 |

### ATAC-seq analysis in patient samples

ATAC-seq primary patient samples reads were mapped against the hg38 reference genome using Bowtie2^91^. ATAC-seq replicates for individual patient samples were pooled. ATAC-seq peaks were called using MACS2. A reference list of peaks or peakome representing all possible ATAC-seq peaks across patient samples was compiled by taking the union of all peaks and merged overlapping peaks. This resulted in a peakome of about 115,457 peaks. Differential analysis was performed using DESeq2^32^ and differentially accessible regions were considered significant if they had an adjusted p-value < 0.01 and a |log2 fold change| >1. Hierarchical clustering was performed on significant differential ATAC-seq peaks (1678) with euclidean distance and complete clustering method with pheatmap R package. Genomic annotation of significant differential ATAC-seq peaks was done using ChIPseeker^92^ with a 3kb window on either side of the TSS, to associate promoter peaks. Bigwigs were obtained for visualization using Deeptools/2.3.3^90^ (parameters bamCoverage --binSize 1 --normalizeUsing RPKM). Heatmaps indicating average ATAC-signal at differentially expressed genes were generated using Deeptools.

### RNA-seq analysis in patient derived cell lines

Paired-end reads were mapped to the hg38 genome using TopHat2^88^ (parameters:–no- coverage-search–no-discordant–no-mixed–b2-very-sensitive–N 1). Counts for genes were obtained using htseq-counts^89^. Differential analysis was performed using DESeq2^32^ and differential genes were considered statistically significant if they had an adjusted p-value less than 0.01 and |log2 FoldChange| >2.

### TARGET RNA-seq analysis of 868 primary patient samples

We downloaded 868 B-ALL patient samples from the TARGET database and samples were analyzed using the sns workflow (https://github.com/igordot/sns) to align and generate counts for genes. Specifically, paired-end reads were mapped to the hg38 genome from GENCODE using STAR aligner^93^ and counts for the 57,316 GENCODE genes (includes coding genes, lncRNAs and pseudogenes) were obtained. Data was normalized using DESeq2^32^.

### Identifying the most likely TF regulators used for network inference

To generate a list of candidate TF regulators, we identified a list of enriched TF motifs at differentially expressed ATAC-seq peaks at promoters (793) previously annotated (**Figure 5e**) in *CRLF2*-High versus Other B-ALL patients using Analysis of motif enrichment (AME^35^) to identify enriched TF motif that are present in the Cis-BP^94^ motif database. AME scores a set of regions with a motif compared to a control sequence. Here we used randomly shuffled input sequences as control sequences and used the one-tailed Fisher’s Exact test to test for motif enrichment. TF motifs with an adjusted p-value< 1e^-13^ were considered enriched TF motifs at promoters with differentially accessible ATAC-seq peaks, which led to a list of 138 TFs. This allows us to limit the number of TF-gene regulatory hypothesis tested.

### Estimating transcription factor activities (TFA)

To generate the prior matrix from the *CRLF2*-high patient cohort, human TF binding motifs (PWMs) were downloaded from the Cis-BP^94^ motif database. Next, we used FIMO^95^ to scan for TF motifs at all ATAC-seq peaks (union of all ATAC-seq peaks in *CRLF2*-High cohort) 20kb upstream of any gene TSS. TF motifs found at each 20kb window per gene with a p-value < 0.0001 were counted.. TF motifs were aggregated across all ATAC-seq peaks for each gene. For each TF motif-gene association, a score, a p-value and an adjusted p-value was computed. Only TF-gene associations that had an adjusted p-value less than 0.01 were retained.

To estimate activity, first DESeq2-normalized TARGET expression data was transposed into matrix ***X***, in which columns are individual patient samples and rows are genes. ***P*** is the TF-gene matrix of known prior regulatory interactions between transcription factors (columns) and genes (rows). ***P****_i,k_* is zero if there is no known regulatory interaction between transcription factor *k* and gene *i. **A*** is the activity matrix, where the columns are the individual samples as in ***X*** and rows are the TFs. We model the expression of gene *i* in individual samples *j* as a linear combination of the activities of each TF, *k*, in condition *j*.

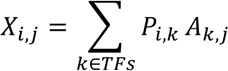

This means that we use the known targets of a transcription factor to derive its activity. The system of linear equations above lacks a unique solution, as there are more equations than unknowns. Hence, to compute a “best fit” (least squares) to the system of linear equations above that lacks a unique solution, we can compute the pseudo-inverse of the prior *P^-1^* multiplied by the expression matrix *X* to approximate *Â*:

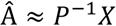

If a transcription factor has no prior targets present in ***P***, we use the expression of that transcription factor as a proxy for its activity.

### B-ALL network inference using (Bayesian Best Subset Regression-Inferelator)

We use a bayesian best-subset regression (BBSR) method, previously described^36^ within the current Inferelator algorithm^25, 26^. At steady state, we model the expression of a gene i in individual cell j as a linear combination of the activities of its TF regulators in individual samples j:

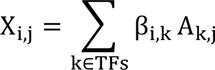

The goal of the network inference procedure is to find a sparse solution for β, in which non-zero entries define both the strength and direction (activation or repression) of a regulatory relationship. This is because we expect a limited number of transcription factors to regulate a particular gene. We select the model with the lowest Bayesian Information Criterion, which adds a penalty term to the training error to account for model complexity and thereby reduce generalization error. After this step, the output is a matrix of inferred regression parameters β, where each entry corresponds to a regulatory relationship between transcription factor k and gene i. Final TF-gene interactions were retained if they had a combined confidence > 0.9 and a precision > 0.3, which resulted in 8,731 regulatory interactions between 135 TFs and 6,645 unique gene targets. Networks were visualized using jp_gene_viz, an interactive interface, designed on iPython. Software is available at https://github.com/simonsfoundation/jp_gene_viz.

### Evaluating Performance of the model

Performance of our model selection within the Inferelator is evaluated by applying a 10- fold cross validation technique using the ATAC-motif prior. The prior matrix was split into 10 equal sized sets. One set is retained as the gold standard to test the model, while the remaining 9 sets are used as the prior. The cross-validation procedure is repeated 10 times, until each of the 10 partitions is used exactly once as the gold standard. To generate a negative control, we randomly shuffle the prior and employ the same 10 fold cross-validation strategy. Next, we compute the area under the precision-recall curve (AUPR) for each of the 10 iterations and compare the results between the prior and the shuffled prior. To test for a significant increase in AUPR values using the ATAC-motif prior compared to a shuffled prior across the 10 iterations, we applied a paired t-test (p-value=4.5e^-5^).

in AUPR values the Overall, AUPR were significantly higher using the ATAC-seq derived motif prior compared to the randomly shuffled prior (**Supplemental Fig 4b**). Thus, the final network was inferred using the ATAC-seq derived motif prior and the number of significant regulatory interactions was limited to TF-gene interactions with combined confidences > 0.9 (high-confidence interactions). This generated a B-ALL specific regulatory network involving 135 TFs (that may be important regulators of *CRLF2*-High gene signature) and 8,731 high confidence patient specific TF-gene interactions (**Fig. 5b**).

### Estimating Transcription Factor Activity in *CRLF2-*High cohort

The B-ALL network was inferred as described above, but to approximate activity of 135 TFs in *CRLF2*-High cohort, we now use the inferred B-ALL network as prior information of TF-gene interaction and gene expression the cohort to estimate activity. Activity values were calculated for each *CRLF2*-High and other B-ALL patient samples. To identify the most likely TFs altered as a result of *CRLF2* overexpression, we used to metrics to define this set of TFs. First, we performed PCA analysis on the activity matrix of 135 TFs. The absolute PC1 values for each TF were computed and ranked to identify the most highly variable TFs that best separate *CRLF2*-High and other B-ALL patient samples, we randomly selected the absolute value of the PC1 score > 0.75 as the most highly variable TFs. Next, we calculated the average TFA for these TFs and a t-test was used to identify TFs with a significant difference in mean estimated TFA between the two conditions (adjusted p-value < 0.001). As a result, we identify 36 TFs that had a significant difference in mean estimated TFA and those with a |PC1| > 0.75.

### Determining the *CRLF2*-associated sub-network affected by *CRLF2* overexpression

Using the inferred B-ALL regulatory interactions, we extract the subset of differentially expressed genes (542) regulated by the 36 significant TFs to derive the *CRLF2*-associated sub- network. Pathway analysis of these genes (Ingenuity Pathway Analysis) resulted in 22 significant pathways (–log_10_(B-H p-value) > 1.3 and a |z-score| >1). Differentially expressed gene targets in the CRLF2-associated sub-network were overlapped with convergent differentially expressed genes in patient derived cell-lines. This resulted in 67 gene targets within the *CRLF2*-associated subnetwork that are robust and convergently differentially expressed across patient and patient derived cell line samples. All sub-networks were visualized using jp_gene_viz (https://github.com/simonsfoundation/jp_gene_viz)

### Validating inferred gene targets with ENCODE CRISPRi and siRNA RNA-seq samples

FOXM1 CRISPRi duplicate RNA-seq samples were downloaded from ENCSR701TVL experiment. Non-targeting control samples were downloaded from ENCSR095PIC experiment. Samples were analyzed using the sns workflow (https://github.com/igordot/sns) to align and generate counts for genes. Specifically, paired-end reads were mapped to the hg38 genome from GENCODE using STAR aligner^93^ and counts for the 57,316 GENCODE genes (includes coding genes, lncRNAs and pseudogenes) were obtained. Differential expression analysis was performed using DESeq2^32^ and genes that had an adjusted p-value < 0.05 and |log2FoldChange|>0.5 were considered significantly altered at the transcriptional levels. This set of genes was overlapped with the number of predicted FOXM1 gene targets to identify what percentage of predicted gene targets could be validated with the CRISPRi dataset. A hyper- geometric test was used to test whether the number of validated gene targets was statistically significant, specifically using phyper in R as follows: phyper(50, 11918, 57316 -11918, 123, lower.tail = FALSE) E2F6 siRNA duplicate RNA-seq samples were downloaded from ENCSR820EGA experiment. Non-targeting control samples were downloaded from ENCSR095PIC experiment. Samples were analyzed using the sns workflow (https://github.com/igordot/sns) to align and generate counts for genes. Specifically, paired-end reads were mapped to the hg38 genome from GENCODE using STAR aligner^93^ and counts for the 57,316 GENCODE genes (includes coding genes, lncRNAs and pseudogenes) were obtained. Differential expression analysis was performed using DESeq2^32^ and genes that had an adjusted p-value < 0.05 and |log2FoldChange|>0.5 were considered significantly altered at the transcriptional levels. This set of genes was overlapped with the number of predicted E2F6 gene targets to identify what percentage of predicted gene targets could be validated with the E2F6 siRNA dataset. A hyper- geometric test was used to test whether the number of validated gene targets was statistically significant, specifically using phyper in R as follows: phyper(57, 3836, 57316 -3836, 272, lower.tail = FALSE)

MAZ siRNA duplicate RNA-seq samples were downloaded from ENCSR129WCZ experiment. Non-targeting control samples were downloaded from ENCSR095PIC experiment. Samples were analyzed using the sns workflow (https://github.com/igordot/sns) to align and generate counts for genes. Specifically, paired-end reads were mapped to the hg38 genome from GENCODE using STAR aligner^93^ and counts for the 57,316 GENCODE genes (includes coding genes, lncRNAs and pseudogenes) were obtained. Differential expression analysis was performed using DESeq2^32^ and genes that had an adjusted p-value < 0.05 and |log2FoldChange|>0.5 were considered significantly altered at the transcriptional levels. This set of genes was overlapped with the number of predicted MAZ gene targets to identify what percentage of predicted gene targets could be validated with the MAZ siRNA dataset. A hyper- geometric test was used to test whether the number of validated gene targets was statistically significant, specifically using phyper in R as follows: phyper(4, 2599, 57316 -2599, 26, lower.tail = FALSE)

### Validating predicted gene targets with ENCODE ChIP-seq samples

Optimal IDR ChIP-seq peaks were downloaded for 6 TFs from ENCODE from the following experiments:

| TF | Experiment | Cells |
| --- | --- | --- |
| **ETS2** | ENCSR596IKD | K562 |
| **TBP** | ENCSR000EHA | K562 |
| **PBX2** | ENCSR263DFP | K562 |
| **BCL11A** | ENCSR000BHA | GM12878 |
| **IRF1** | ENCSR000EGU | K562 |
| **EGR1** | ENCSR211LTF | K562 |

Genomic annotation of optimal IDR peaks was done using ChIPseeker^92^ with a 20kb window upstream of the TSS, to associate peaks to genes. The same threshold was used to generate TF-gene interactions for the prior matrix used for network inference. Genes with at least one optimal IDR peak were overlapped with predicted gene targets for each TF from the B-ALL network. This resulted in the number of predicted gene targets supported by ChIP-seq binding of each TF of interest. Hyper-geometric tests were applied to test whether the number of validated gene targets was statistically significant for each TF.

## Acknowledgements

The authors thank all members of the Skok, Bonneau, and Carroll labs for discussions. The Children’s Oncology Group for providing us with the patient samples and metadata. New York School of Medicine High Performance (HPC) for technical support. Adriana Heguy and the Genome Technology Center (GTC) core for sequencing efforts. The TARGET database.

## Author contributions

B.C, S.B, R.B and J.S designed and managed the project. B.C, P.L, and N.E collected samples and performed experiments. S.N and W.C provided experiments. S.B performed data analyses. D.C, C.S, R.R, and A.W aided in analysis design. B.C, R.B, and J.S secured funding. S.B and J.S wrote the manuscript. R.B and J.S revised the manuscript.

